# Deep mutational scanning and machine learning uncover antimicrobial peptide features driving membrane selectivity

**DOI:** 10.1101/2023.07.28.551017

**Authors:** Justin R. Randall, Luiz C. Vieira, Claus O. Wilke, Bryan W. Davies

## Abstract

Antimicrobial peptides commonly act by disrupting bacterial membranes, but also frequently damage mammalian membranes. Deciphering the rules governing membrane selectivity is critical to understanding their function and enabling their therapeutic use. Past attempts to decipher these rules have failed because they cannot interrogate adequate peptide sequence variation. To overcome this problem, we develop deep mutational surface localized antimicrobial display (dmSLAY), which reveals comprehensive positional residue importance and flexibility across an antimicrobial peptide sequence. We apply dmSLAY to Protegrin-1, a potent yet toxic antimicrobial peptide, and identify thousands of sequence variants that positively or negatively influence its antibacterial activity. Further analysis reveals that avoiding large aromatic residues and eliminating disulfide bound cysteine pairs while maintaining membrane bound secondary structure greatly improves Protegrin-1 bacterial specificity. Moreover, dmSLAY datasets enable machine learning to expand our analysis to include over 5.7 million sequence variants and reveal full Protegrin-1 mutational profiles driving either bacterial or mammalian membrane specificity. Our results describe an innovative, high-throughput approach for elucidating antimicrobial peptide sequence-structure-function relationships which can inform synthetic peptide-based drug design.

## Introduction

Antimicrobial peptides (AMPs), including host defense peptides, are an evolutionarily conserved group of natural peptides which fight pathogens through direct antimicrobial activity and/or by modulating the host immune response during infection^1^. AMPs generally kill bacteria directly through cell membrane disruption, but many show difficulties differentiating bacterial and mammalian cell membranes. This effect has been exploited in some cases where AMPs have been explored as anticancer agents. Some aspects of membrane selectivity have been described for alpha-helical AMPs^2^. However, this information is limited, and the sequence-activity relationship governing AMP membrane selectivity remains largely unclear, which hinders their potential for therapeutic use.

The most thorough means of understanding AMP sequence-activity relationships is through mutational analysis. However, current methods for exploration of antimicrobial peptide residue importance and flexibility are slow and generally limited to small numbers of variants. One still popular method is to perform alanine scans by changing each position individually to alanine and measuring its effect on structure or activity. Such analog methods limit the number and diversity of mutations which can be realistically examined.

Recently, we developed a new high-throughput method called surface localized antimicrobial display (SLAY), which has made it possible to assess the antibacterial potential of hundreds of thousands of peptide sequences simultaneously^11,12^. The simultaneous development of deep mutational scanning of protein sequences now allows for large scale assessment of up to millions of mutations on a measurable activity of interest^13^. Here we integrate these two approaches to create deep mutational SLAY scanning (dmSLAY), which can assess how thousands of sequence variations impact AMP activity simultaneously. We apply the unprecedent power of dmSLAY to investigate the AMP Protegrin-1. Protegrin-1 (PG-1), originally discovered in porcine leukocytes, is a member of the rare group of AMPs with β-hairpin structure cyclized via disulfide bonds (β-AMPs). β-AMPs kill primarily via membrane lysis but show a wide range of antibacterial potency and mammalian toxicity which remains mostly unresolved^3^. Protegrin-1 (PG-1), demonstrates potent and broad-spectrum bacterial membrane disruption^4,5^, but it also shows high levels of hemolysis and cytotoxicity^6,7^. Still, PG-1 has been successfully used as a scaffold by academics and biotechnology companies to design clinically relevant antibiotics such as Murepavadin^8,9^. Though promising, phase III clinical trials with Murepavadin were ultimately terminated due to toxicity concerns^10^. Identifying the sequence features associated with the toxic side effects of similar β-AMPs could help fully realize the true clinical potential of this class. Applying dmSLAY to PG-1 uncovered rules governing its membrane selectivity and enabled us to train machine learning algorithms to digest millions of additional sequence variants and reveal important sequence-structure-function relationships influencing PG-1 bacterial and mammalian membrane selectivity. Our results establish dmSLAY as a new method to gain unprecedented insight into AMP sequence-activity relationships and reveal knowledge that could be used to improve rational optimization of β-AMPs to reduce their toxicity and advance their potential for clinical use.

## Results

### Deep mutational scanning of the Protegrin-1 sequence

We began our study of PG-1 by performing a traditional alanine scan in an effort to help elucidate residues which might be important for antimicrobial activity (**Fig. S1A**); This scan however proved rather ineffective, since the only residue changes causing more than a two-fold change in activity were R4A, Y7A, and R19A, which were four-fold less potent by measurement of minimum inhibitory concentration (MIC).

To overcome the shortfalls of alanine-type scans and more efficiently elucidate PG-1 residue importance and flexibility, we developed deep mutational surface localized antibacterial display (dmSLAY). To accomplish this we first designed a genetically encoded deep mutational PG-1 variant library that was compatible with our antibacterial display approach. This library encoded 7 to 12 possible single residue mutations at each residue position and 1 to 9 residue changes per unique variant (**Fig. S1B**). In total this library consists of 7,105 unique PG-1 sequences, 183 of which are single residue variants. This is a massive increase in diversity over the 18 single mutants provided by traditional alanine scanning.

We next cloned the PG-1 variant library into an inducible surface display plasmid and transformed the library into *E. coli* W3110 for high-throughput examination of antimicrobial potential using SLAY. SLAY exploits a bacterium’s ability to both produce and simultaneously screen large peptide libraries for antibacterial potential. Each library is displayed on the outer membrane of bacteria by a tether fused to a fragment of OmpA. If a peptide is antibacterial, the bacterium expressing it is eliminated from the population only when expression is induced. Next-generation sequencing can quantify plasmids isolated from induced and uninduced cultures to generate a log_2_-fold change in reads predicting each peptide sequence’s antimicrobial potential (**Fig. 1A**).

**Figure 1:**
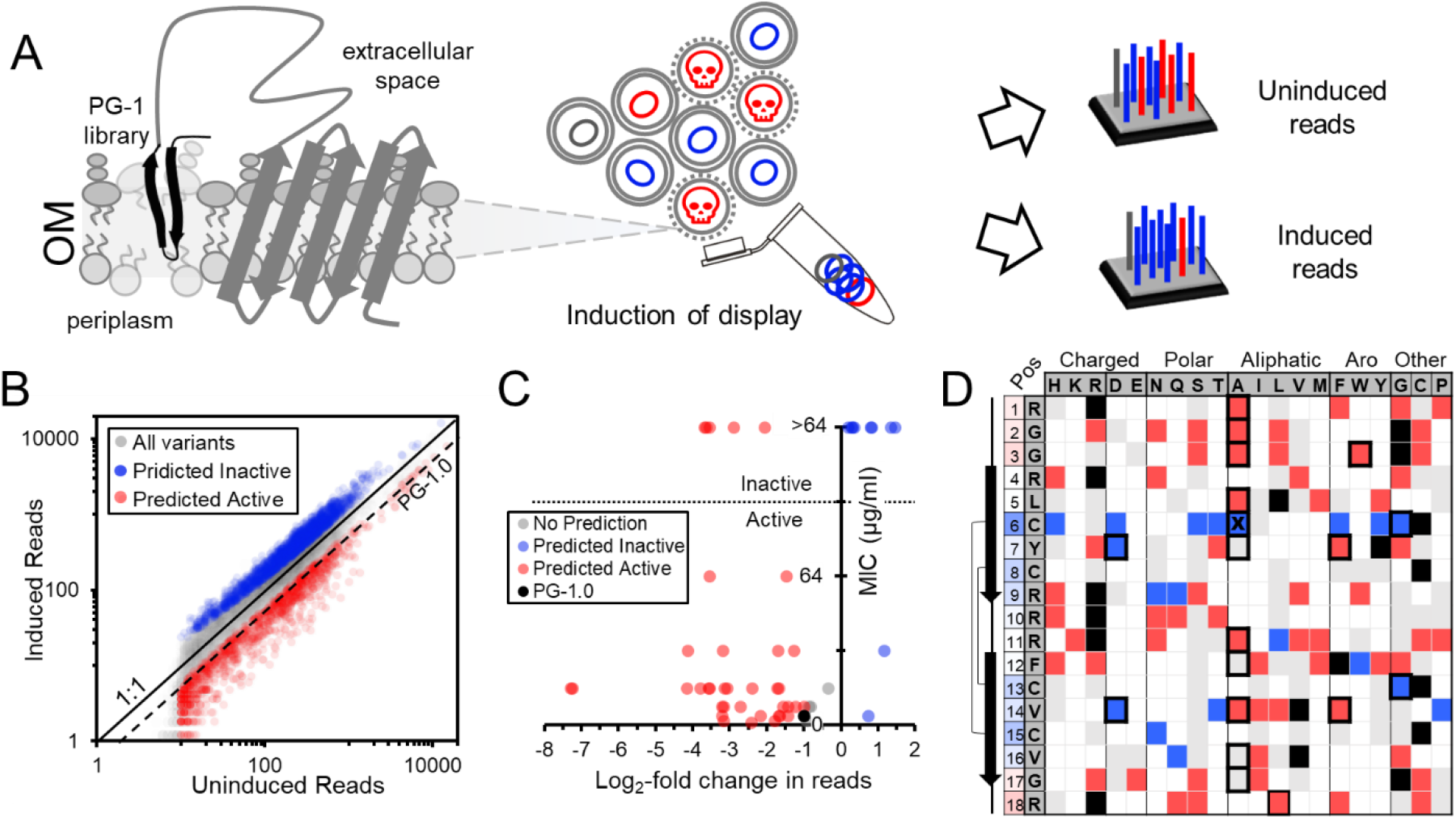
Protegrin-1 deep mutational SLAY predicts residue importance and flexibility. A) Surface localized antimicrobial display expresses an OmpA fusion protein tethering the PG-1 library to the outer membrane (OM). Induction of display results in cell death for variants maintaining antimicrobial activity (red-active, blue-inactive). change in reads between induced and uninduced cultures can predict antimicrobial potential. B) Scatter plot of induced versus uninduced reads for the entire PG-1 library on log scales. 1:1 and the native PG-1 sequence (PG-1.0) read ratios are shown. C) Scatter plot of MIC for a subset of 52 variants in the PG-1 library versus the log_2_-fold change in reads. D) Table charting dmSLAY predictions for all single PG-1 variants in the library. The native PG-1 sequence and position in columns one and two, amino acid change at each position going across the top categorized by side chain. PG-1 secondary structure and disulfide bonds are diagramed to the left. Position column is color coded by average log_2_-fold change. Bold boxed cells were evaluated *in vitro*. An X marks an incorrect dmSLAY prediction.

A SLAY screen run on our PG-1 variant library identified 1,940 likely inactive variants with a significant log_2_-fold change (p < 0.05) in reads greater than zero (**Fig. 1B**, blue). As expected, the native PG-1 sequence (PG-1.0) had a negative log_2_-fold change in reads. Using the PG-1.0 log_2_-fold change as a benchmark, another 1,203 variants were predicted to retain their antimicrobial activity (log_2_-fold change < -1, p < 0.05) (**Fig. 1B**, red). No prediction was made for variants with a log_2_-fold score between 0 and -1 since values in this range frequently lacked significance (p value > 0.05).

### dmSLAY accurately predicts Protegrin-1 residue importance and flexibility

To investigate the accuracy of dmSLAY predictions, we synthesized 40 PG-1 variants spanning a range of log_2_-fold change scores and measured their MIC against *E. coli* W3110 in MH media (**Table S1**). We also included data from eleven alanine scan variants already examined. Only variants with an MIC less than or equal to 64 μg/ml were deemed to be antibacterially active. The log_2_-fold change score did not correlate directly with antibacterial potency, but a negative or positive score was 86% accurate at predicting whether a variant retained or lost antimicrobial activity (> or ≤ 64 µg/ml) (**Fig. 1C**). This included 16/17 accurate predictions for single residue PG-1 variants examined *in vitro* (**Fig. 1D**, bold bordered cells), and indicated dmSLAY was highly accurate at predicting if a PG-1 variant retained or lost antibacterial activity.

PG-1 has an antiparallel β-sheet structure consisting of two sheet regions with interstrand disulfide bonds, a loop region in between them, and two tail regions near either terminus^14^ (**Fig. 1D, Fig, S2A**). The predicted antibacterial activity for each single variant from dmSLAY is cataloged in a chart and the average log_2_-fold change for each individual PG-1 position (1-18) across all possible residue changes is colored by varying degrees of blue (inflexible) or red (flexible) (**Fig. 1D**, far left column). Residues in the loop and tail regions of PG-1 were generally the most flexible while residues in the sheet regions were predicted to be more important for antibacterial activity. Cysteines involved in disulfide bonds were the least flexible residues, suggesting disulfide bonds maybe especially important for antimicrobial activity. It is important to note that many residue positions had amino acid changes predicted to either retain activity or lose activity, highlighting the importance of evaluating more than just one residue change at each position.

To evaluate mutations observed in PG-1 variants with multiple residue changes, a sequence logo plot was generated for the 50 variants with the lowest and highest dmSLAY log_2_-fold change scores (**Fig. S2**). Multiple residue variants predicted to lose activity often had mutations within the β-sheet regions, especially the cysteine residues, consistent with observations from single variants. Variants retaining activity mostly contained changes in the loop and tail regions, but some contained changes to sheet residues with similar side chain amino acids.

### Changes in secondary structure correlate with Protegrin-1 lytic activity

With our unprecedent evaluation of residue importance in PG-1, we next looked to elucidate relationships between PG-1 sequence, structure, and function. To do this we used circular dichroism spectroscopy (CD) to examine changes in secondary structure for the 40 PG-1 variants selected above (**Table S1**). Many AMPs, including PG-1, show an unstructured CD spectrum in buffer alone but fold into alpha-helical or β-sheet structure upon interaction with membranes or membrane mimics^15^. With this in mind, CD spectra were measured for PG-1.0 and all 40 variants with and without the presence of lipopolysaccharide (LPS), to mimic the bacterial outer membrane. As expected, native PG-1, and almost all its variants switched from unstructured (minimum at ∼200 nm) to β-sheet (minimum at ∼220 nm, maxima at ∼195) upon addition of LPS^16^ (**Fig. 2AB**). To determine how secondary structure may impact antimicrobial activity, we color coded spectra by MIC. Without LPS we saw no discernable correlation between CD spectra and MIC (**Fig. 2A**); however, upon addition of LPS we found that variants with a spectrum more representative of native PG-1 (PG-1.0) were generally more potent than those with larger changes in spectra (**Fig. 2B**). This suggests that the native secondary structure of PG-1 upon interaction with cell membranes is important for its antibacterial activity.

**Figure 2.**
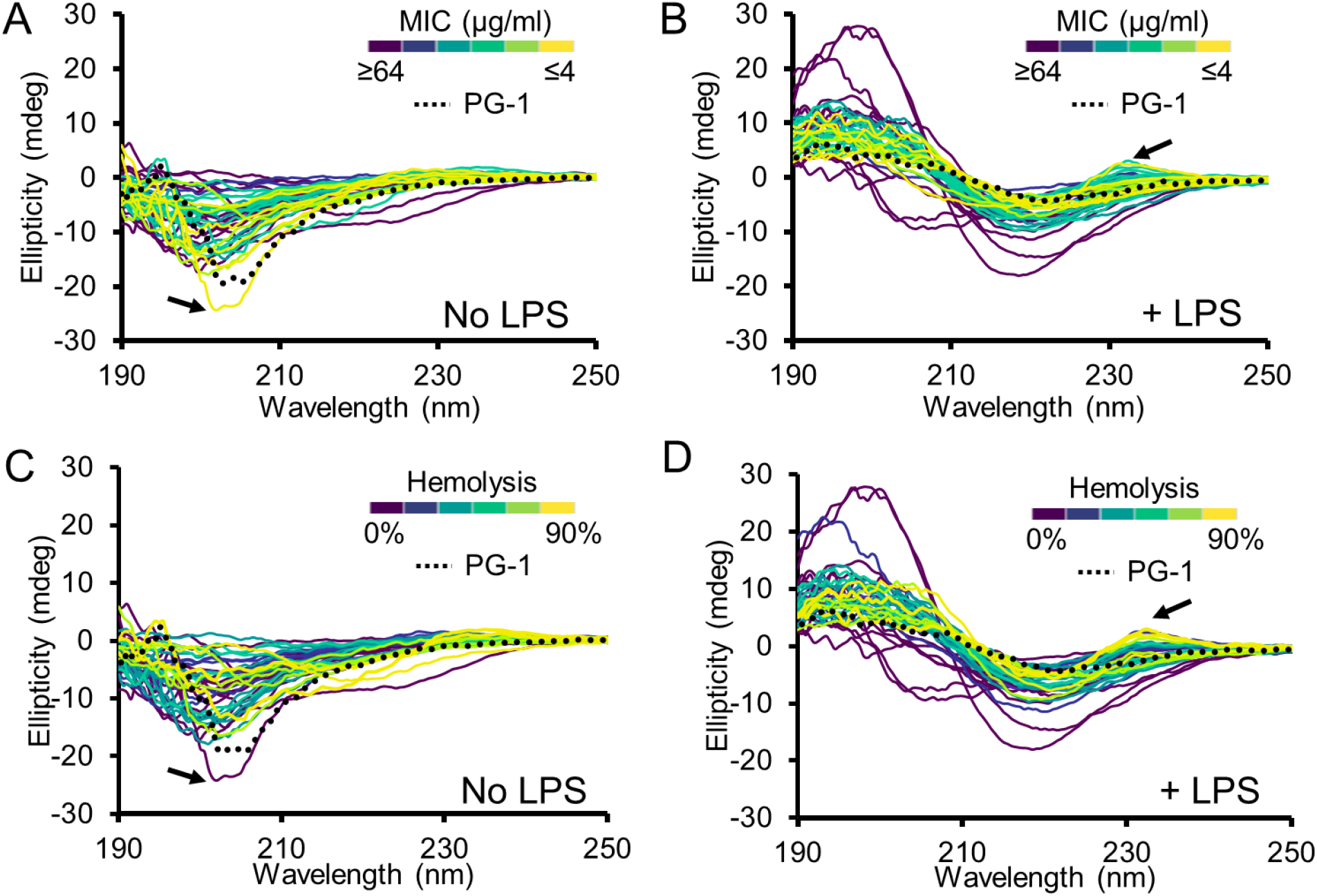
Changes in secondary structure correlate with Protegrin-1 lytic activity. Circular dichroism spectra select PG-1 variants from the dmSLAY scan color coded by minimum inhibitory concentration (A, B) or percent hemolysis (C, D). The native PG-1 sequence (PG-1.0) spectrum is shown as a dotted line. Arrows indicate different trends in cell membrane lysis for PG-1.37 (A, C) or tryptophan containing variants with a maximum near 230 nm. All spectra are the average of technical triplicate.

Since PG-1 kills bacteria by cell membrane lysis, we questioned whether this phenomenon was also true for lysis of mammalian membranes; therefore, we measured the hemolytic activity of each variant (**Table S1**) and color-coded spectra by the percent hemolysis observed from treatment with 128 μg/ml peptide over three hours (**Fig. 2CD**). Again, we observed no correlation in percent hemolysis and CD spectra in the absence of LPS, but upon its addition we saw a similar correlation observed for MIC where hemolytic variants retained secondary structure close to PG-1.0 upon membrane interaction. One interesting difference was increased hemolytic spectrum with a maximum at 230 nm (**Fig. 2BD**, arrows). This peak is found only in peptides containing tryptophan and is consistent with previous research showing tryptophan residues generally increase mammalian cell toxicity^17–19^.

### Membrane selectivity is influenced by absence of large aromatics and loss of cysteine pairs

We noticed that some PG-1 variants appeared to have lost their hemolytic activity but remained strongly antibacterial, suggesting increased membrane selectivity (**Fig. 2AC**, arrows). To better identify these selective variants, we generated a selectivity score for each variant tested by multiplying the MIC and the %hemolysis. Variants with a lower selectivity score are more bacterially selective. We compared the selectivity score for all active PG-1 variants to PG-1.0 to identify those with improved membrane selectivity, resulting in five such variants (**Fig. 3A**). We again color-coded CD spectra in the presence of LPS, this time by selectivity score, and found that spectra with a maximum at 230 were generally less selective with the exception of PG-1.34 (**Fig. 3B, Table S1**), but no other obvious correlations were observed. This is again consistent with peptides containing tryptophan residues having higher mammalian toxicity without impacting antibacterial activity. β-hairpin peptides are prone to aggregation which could also relate to their cellular toxicity. We tested if toxicity correlated with the propensity of PG-1 variants to self-aggregate and found an exponential relationship (R^2 = 0.63) where highly hemolytic variants (>75%) were also highly aggregative and less hemolytic variants (< 30%) were generally non-aggregative (**Fig. 3C**); however, there was a wider range of aggregation in variants with hemolytic activities between 30% and 75% and the relationship did not fully explain differences observed in selectivity score.

**Figure 3.**
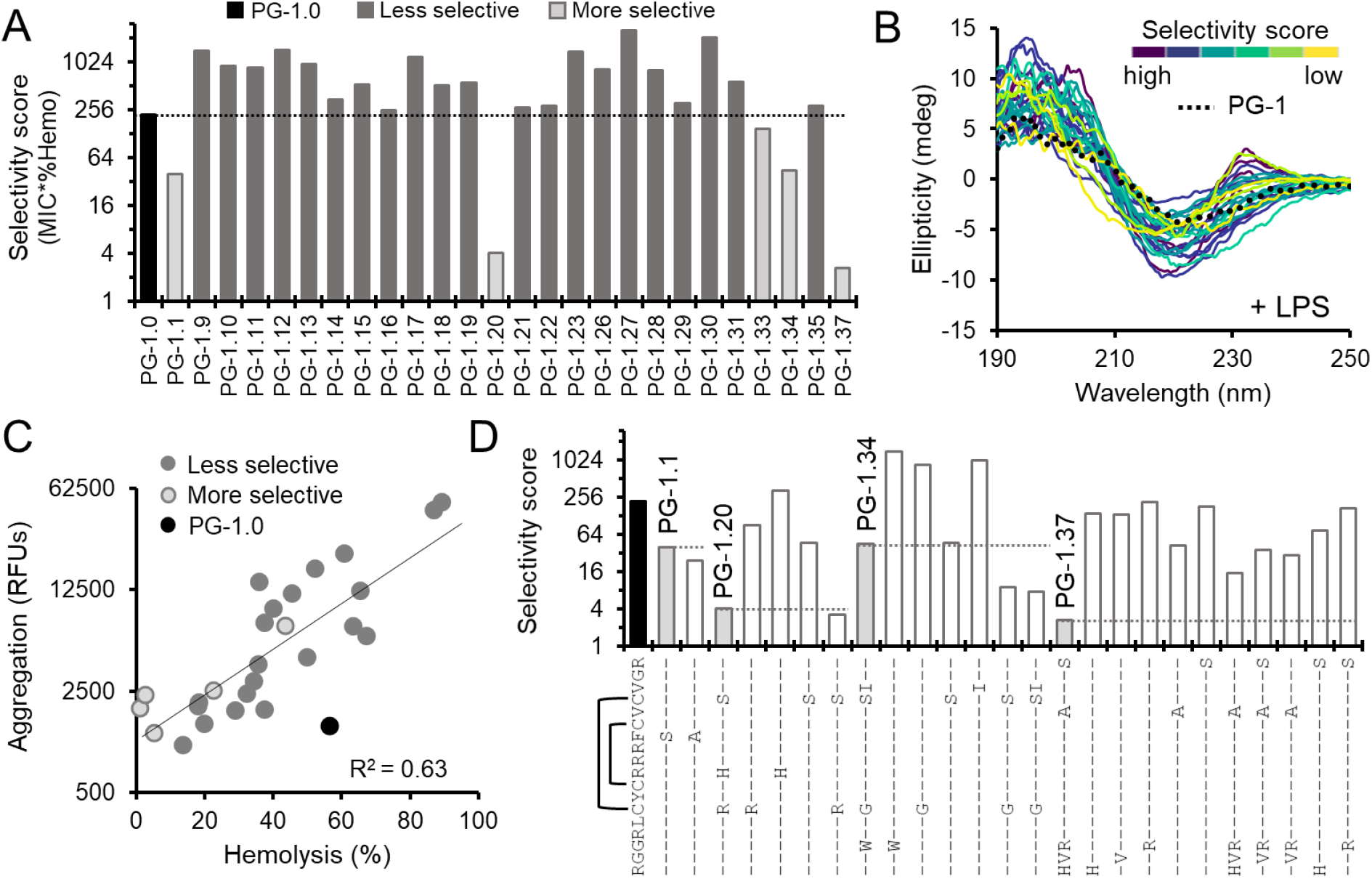
Membrane selectivity is influenced by aromatics and loss of cysteine pairs. A) Bar graph comparing selectivity score (median minimum inhibitory concentration * average percent hemolysis) for PG-1 variants confirmed to be antibacterial on a log scale. B) Circular dichroism spectra of same PG-1 variants color coded by selectivity score. Spectra are the average of technical triplicate. C) Scatter plot of PG-1 variant aggregation in relative fluorescent units (RFUs) on a log scale versus linear percent hemolysis. Each data point is the mean of triplicate reactions, trendline represents a linear fit of log transformed aggregation data versus hemolysis with R^2^ = 0.63. D) Bar chart of selectivity score for a subset of mutations found in the four most selective PG-1 variants on a log scale. Brackets indicate where disulfide bonds are present in the native PG-1 sequence.

Since changes in structure and self-aggregation did not explain differences observed in membrane selectivity, we broke down the sequence changes found in the four most selective variants to explore whether specific mutations influence selectivity (**Fig. 3D**). We found that mutation of phenylalanine at position 12 to either a serine or alanine accounted for the selectivity observed for PG-1.1, suggesting that phenylalanine itself may have a negative effect on selectivity. This supports our overall observation that large aromatics in PG-1 promote hemolysis and decreased selectivity. Unexpectedly, we identified cysteine mutants that retained antibacterial activity and improved their selectivity. Corresponding C6R and C15S mutations accounted for the increased selectivity of PG-1.20, whereas C15S alone matched PG-1.34’s selectivity score; however, a corresponding mutation of C6G in PG-1.34 further reduced its selectivity score. This data suggests that removal of the disulfide bond between position 6 and 15 to certain amino acids which don’t strongly impacting PG-1 structure, may improve bacterial membrane selectivity. While most individual mutations of cysteines residues are not well tolerated (**Fig. 1D**), insights gained through dmSLAY show certain exceptions to this rule can improve selectively. Removal of the G3W mutation in PG-1.34 also caused a decrease in selectivity score, further supporting that tryptophan generally increases mammalian toxicity. PG-1.37 contains no changes to positions 6, 12, or 15 and its selectivity could only be recapitulated by the presence of all five mutations, suggesting influences on selectivity for PG-1 variants with many mutations may be more complicated than simple removal of disulfide bonds and/or large aromatic residues; however, these variants can be identified using deep mutational scanning.

We noticed that all five more selective variants contained residue changes to serine and/or histidine (**Table S1**). To see if serine/histidine mutations have any general impact on selectivity, we synthesized ten additional PG-1 variants containing either a serine or histidine residue change with a significant negative log_2_-fold change from dmSLAY (**Fig. S3A**). We used single variant predictions from dmSLAY to aid our selection (**Fig. 1D**). 6/10 variants from this group had improved selectivity compared to native PG-1, but these improvements were more modest than those observed with PG-1.1, PG-1.20, and PG-1.37 (**Fig. S3B**). Five of the six more selective variants contained mutations to the phenylalanine at position 12 or compatible mutations of two cysteines involved in a disulfide bond, suggesting again that removal of phenylalanine and/or compatible mutations to cysteines involved in a disulfide bond likely improves bacterial membrane selectivity (**Fig. S3**). PG-1.57 (V14A, R18H) and PG-1.37 were the only more selective PG-1 variants which did not contain these specific mutations.

### Protegrin-1 variants demonstrate strong specificity for bacterial membranes

PG-1 kills bacteria via disruption of the outer and inner cell membrane. To ensure that our selective variants were killing via the same mechanism of action, we performed propidium iodide (PI) uptake assays. PI is a membrane impermeable dye which becomes fluorescent upon interaction with DNA; therefore, both the outer and inner membrane must be permeabilized for PI to access the cytoplasm and give a strong fluorescent signal. *E. coli* W3110 cells treated with serially diluted PG-1.0 for 30 minutes showed strong uptake of PI, as expected (**Fig. 4A**). Similarly, our selective variants also showed strong PI uptake, suggesting they likely work via the same membrane lytic mechanism of action. Membrane lytic AMPs generally cause rapid cell death within 30 minutes of treatment. Kill curves performed with PG-1.0 and selective variants at concentrations four-fold their MIC show near complete *E. coli* W3110 cell death within 30 minutes, further supporting these variants kill via the same membrane lytic mechanism of action (**Fig. 4B**).

**Figure 4.**
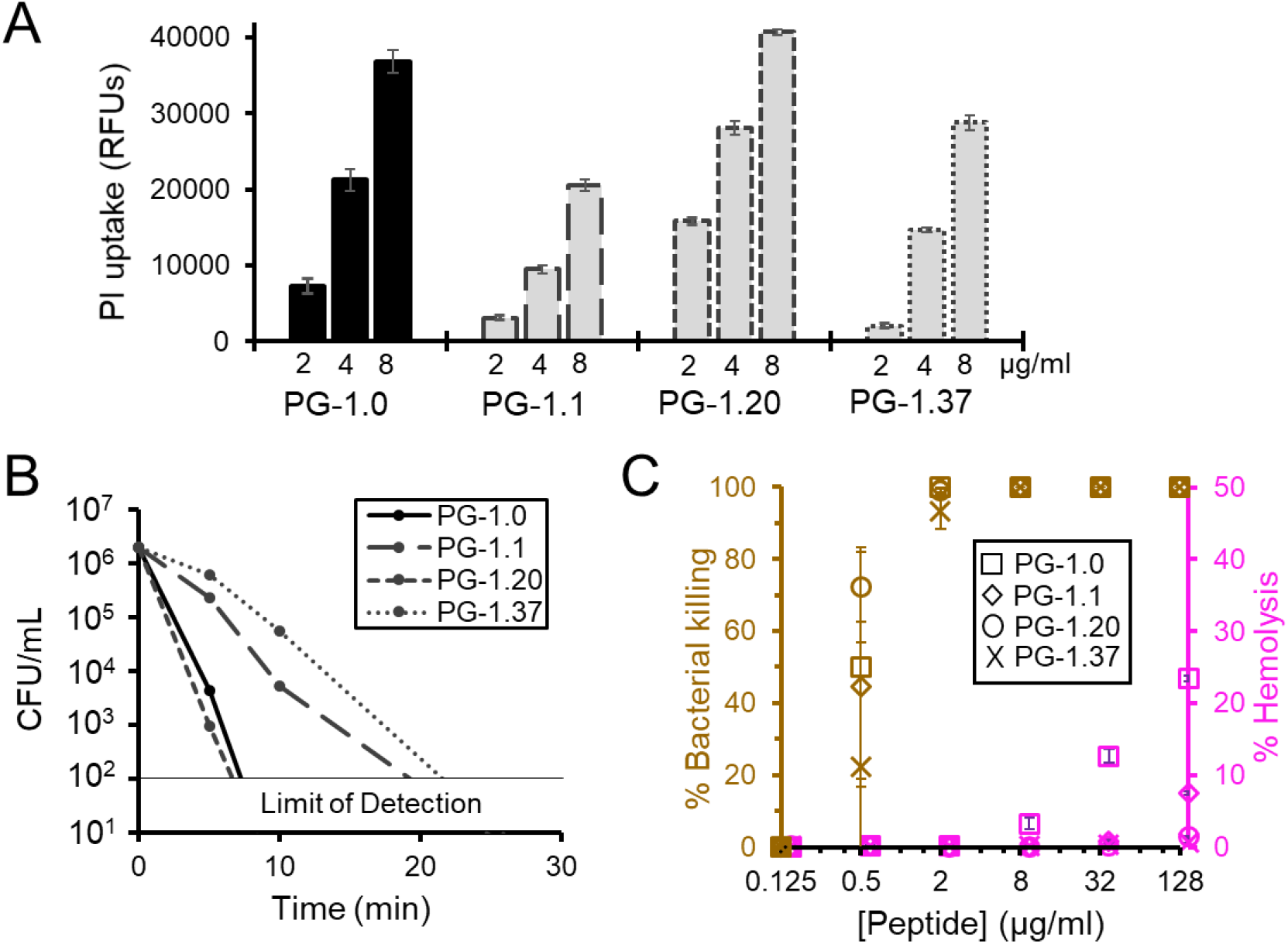
Protegrin-1 variants demonstrate strong specificity for bacterial membranes. A) Bar chart of propidium iodide (PI) uptake measured in relative fluorescent units (RFUs) for *E. coli* cells treated with the indicated concentration of each PG-1 variant. B) Kill curve showing colony forming units (CFUs) present over time for cultures treated with 8 µg/ml peptide. C) Percentage of bacterial killing and hemolysis observed in co-culture of *E. coli* and human red blood cells treated with increasing concentrations of PG-1 variants after one hour of treatment. All data points in this figure are the mean of triplicate reactions. Error bars represent one standard deviation.

To examine in detail PG-1 variant bacterial membrane specificity, we simultaneously treated *E. coli* W3110 cells and human red blood cells with increasing concentrations of each variant at the same time. After one hour of treatment we measured both the percentage of bacterial killing and the percent hemolysis occurring at each concentration (**Fig. 4C**). Bacterial killing was first observed at 0.5 μg/ml for each variant. PG-1.20 and 1.37 showed no significant hemolytic activity, even at the highest concentration tested, while PG-1.1 and PG-1.0 began to noticeably lyse red blood cells at 128 and 8 μg/ml respectively, confirming previous observations when treated individually, suggesting these variants are more toxic to red blood cells than PG-1.20 and PG-1.37. When you compare each variant’s bacterial killing with and without the addition of red blood cells, we see no significant reduction in bacterial killing upon addition of 100-fold more red blood cells. This equates to greater than 1000-fold more mammalian cell membrane surface area when considering differences in cell size (**Fig. S4**). The inability of overwhelming amounts of mammalian cell membrane to competitively inhibit antibacterial activity suggests our variants contain greater than 1000-fold more specificity for bacterial membrane compared to red blood cell membrane. Surprisingly, this also remained true for PG-1.0 despite the hemolysis observed at higher concentrations.

### Machine learning identifies mutational profiles promoting membrane specificity

dmSLAY revealed several unexpected PG-1 variants with increased selectivity; however, this data is not sufficient for rational design of new PG-1 variants with desired qualities. Deep mutational scanning predicted the antibacterial potential of over 4000 PG-1 sequence variants covering ∼47% of possible single residue changes, but it only assessed less than 0.1% of possible variants containing two or three residue changes. As seen with the selective PG-1 variants examined thus far, combinations of mutations can have a much greater effect on bacterial membrane selectivity then single mutations alone. To assess the full spectrum of possible one, two, and three residue variants impacting function (> 5.7 million) we trained three different machine learning models with our new biochemical dataset. These three algorithms focused on predicting PG-1 variant antibacterial potency (**Fig. S5A**), toxicity (**Fig. S5B**), and bacterial membrane selectivity (**Fig. S5C**), respectively. We trained each algorithm on 80% of our data and used the final 20% to validate the predictions made by the final models (**Fig. S5**, top panels). We then used these three models to identify the mutational profiles which help maintain PG-1 antibacterial potency (MIC ≤ 8 µg/ml), decrease mammalian toxicity (hemolysis < 2%), or increase bacterial specificity (log_10_-selectivity score < 0.5) individually. Heat maps of these mutational profiles generally show a lack of cysteine addition and increased mutation of the second beta-sheet in PG-1 variants retaining antibacterial activity. Changes are common in β-sheet residues for less hemolytic variants with exclusion of tryptophan and cysteine. More membrane selective PG-1 variants show mutation of β-sheet residues to charged or polar amino acids while lacking addition of all aliphatic and aromatic residues, except alanine (**Fig. S5**, bottom panels). Though these models provide detailed information regarding how PG-1 sequence impacts certain biochemical qualities, individually they do not select solely for highly potent variants specifically targeting only bacterial or mammalian membranes.

To find highly potent PG-1 variants which are non-hemolytic, we combined all three models and identified 17,348 unique sequences passing all three cut offs (**Fig. 5A**). To examine the success of modeling in identifying more bacterially selective variants, we chose thirty-six variants from of the 17,348 and measured their MIC and percent hemolysis *in vitro* (**Table S2**). We then used this information to calculate their selectivity score and compared these attributes to see if they were more bacterially selective than PG-1.0 and if they met the cutoffs used from machine learning predictions (**Table S3**). From this group of thirty-six variants, 81% had an improved selectivity-score compared to PG-1.0 and the average log_2_-fold change in selectivity score was -3.80. Additionally, 25% had an MIC ≤ 8 µg/ml, 89% had under 2% hemolysis, and 17% had a log_10_-selectivity score less than 0.5. For comparison, we contrasted these results to those obtained for the 31 dmSLAY active variants characterized during initial analysis of dmSLAY predictions (**Fig. 1C**, red). Here, only 16 percent were more selective than PG-1.0 and the average log_2_-fold change in selectivity score was 1.68. Also, 29% had an MIC less than 8 µg/ml, 6% had under 2% hemolysis, and only 3% had a log_10_-selectivity score less than 0.5. This means that our machine learning predictions vastly outperformed dmSLAY at identifying membrane selective and non-toxic PG-1 variants while being nearly as successful as dmSLAY at predicting antibacterial potency.

**Figure 5.**
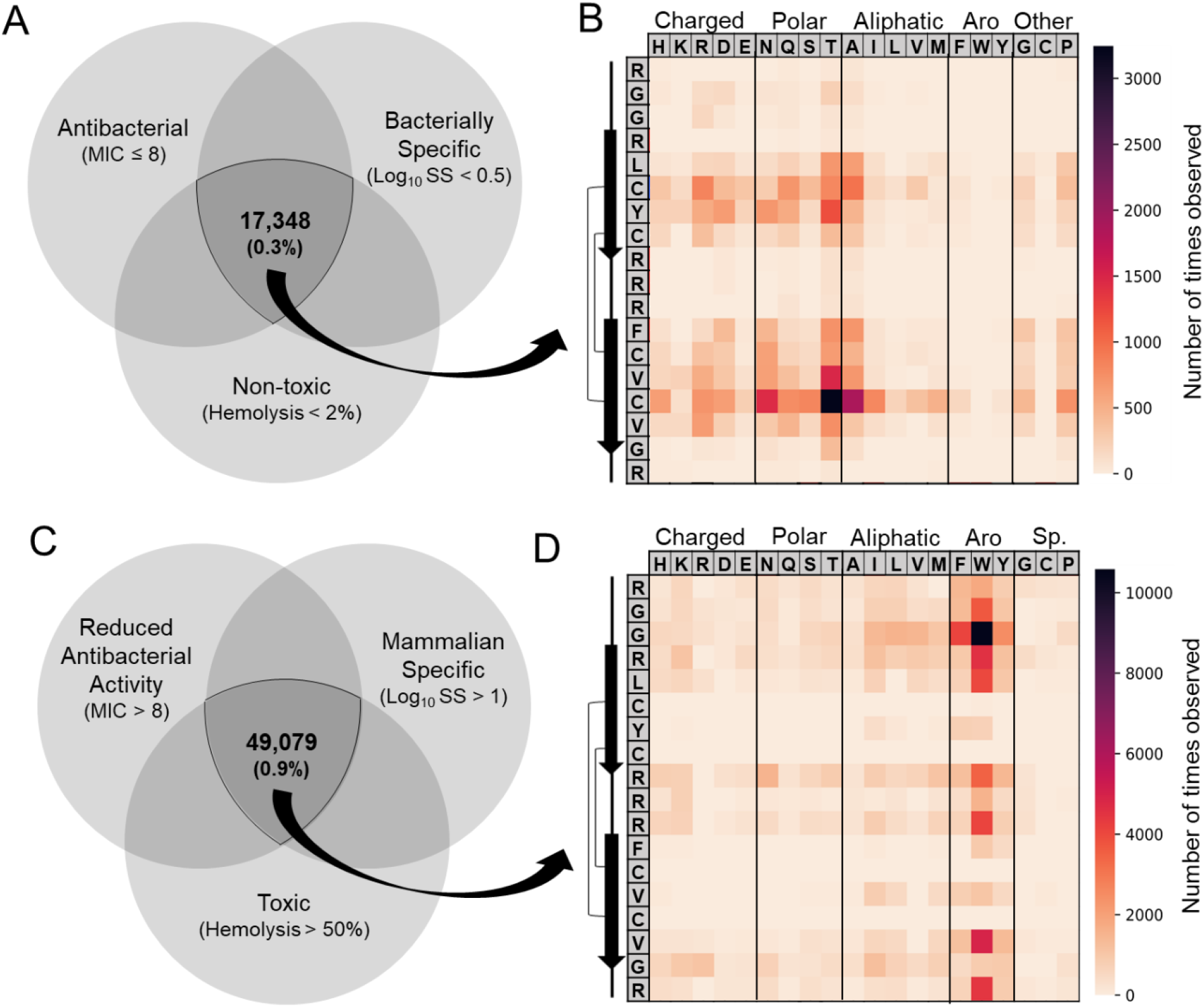
Machine learning identifies mutational profiles promoting membrane specificity. A) Venn diagram showing the three machine learning models trained on PG-1 variant data and the cut off used to identify the 17,348 most bacterially specific (A) or mammalian specific (C) from over 5.7 million one, two and three residue variants. Mutational heat maps charting the number of times each mutation is observed within the bacterially selective group (B) or mammalian selective group (D). The native PG-1 sequence is shown going down the far left column and specific residue change across the top categorized by side chain. Native PG-1 secondary structure is diagramed to the left.

To visualize the mutational profile present in bacterially selective PG-1 variants we generated a heat map showing the number of times each mutation was observed within the bacterially selective group (**Fig. 5B**). Here, hydrophobic residues were changed within the β-sheet regions, especially at C15. These changes were often to polar or charged rather than other hydrophobic or aromatic residues with one notable exception being alanine, possibly due to its overrepresentation within the training dataset. Many of the most common residue changes observed did not exist in the training dataset at all, including C15T which was the most frequent change overall. For comparison, we also used our models to select for mammalian specific variants by adjusting the cutoffs used for each model, identifying 49,079 more mammalian specific sequences. We observed an opposite trend where hydrophobic or aromatic residues, especially tryptophan, replace mostly cationic residues in the loop and tail regions (**Fig. 5CD**). Together, this suggests that reducing hydrophobicity within the β-sheet regions of PG-1 generally improves bacterial membrane specificity while adding hydrophobicity, especially with tryptophan, and in the tail and loop regions while decreasing overall charge improves mammalian membrane specificity. For a full list of variant sequences identified and their corresponding machine learning predictions for each model see Data file 1.

Together, this data demonstrates machine learning is far more successful at identifying PG-1 variants with improved bacterial selectivity than random expectation, while enabling the rapid examination of millions of sequence variants. The combination of dmSLAY variant characterization paired with machine learning is a massive improvement over low-throughput analog methods like alanine scans and other methods of peptide drug development which are generally limited to less than 10,000 short peptides and are generally inaccessible to most research labs^20–22^.

## Discussion

With the critical need for new antibiotics, more and more antimicrobial peptides have begun to enter the clinical pipeline^1^. Mammalian cell toxicity continues to be a major barrier to their success, mostly due to our lack of knowledge regarding sequence-structure-function relationships influencing bacterial membrane specificity. This is largely driven by a lack of data, contradictory evidence, and inconsistencies in sequence factors dictating AMP selectivity^23^. Deep mutational SLAY analysis offers a new high-throughput method of quickly determining positional residue importance and flexibility, while also predicting how antimicrobial activity is influenced by multiple simultaneous sequence changes (**Fig. 1**). The successful application of dmSLAY to PG-1, provides a compelling template for analysis of a wide range of AMP sequences and structures. This type of in-depth analysis across diverse unrelated sequences may help to reveal unifying AMP sequence-structure-activity relationships as well as highlight important differences unique to specific structural classes.

Our structural analysis of PG-1 variants revealed a strong correlation between secondary structure induced by membrane mimics and both bacterial and mammalian membrane lysis (**Fig. 2**). Previous studies have shown a similar correlation between changes in PG-1 secondary structure and cytotoxicity^7^ and correlations between antibacterial activity and hemolysis^24^; however, attempts to identify characteristics responsible for differentiation of these two activities have been generally unsuccessful^6^. Cysteines have been previously identified as important for retention of PG-1 antibacterial activity^24–26^. Our dmSLAY results also support the importance of cysteine in maintaining antibacterial activity for single mutants; however, our work shows that specific corresponding mutations of cysteines involved in disulfide bonds which do not compromise native secondary structure can greatly reduce hemolysis and do so without impacting antibacterial potency (**Fig. 3**). These results demonstrate the importance of examining a more diverse set of sequence changes rather than just single point mutations when assessing AMP residue importance and flexibility.

While dmSLAY offers unprecedented screening scale of AMPs, it can still only evaluate a small number of sequence variants compared to the total number of combinatorial possibilities. Here we amplified our exploration of the PG-1 sequence space by leveraging machine learning. This allowed the virtual evaluation of 5.7 million PG-1 variants for antimicrobial activity, toxicity, and membrane selectivity. Machine learning methods are often trained on very large datasets. Remarkably, we developed highly accurate models of PG-1 activity using relatively few (<100) input sequences. Our modeling provided further insight into PG-1 sequence-activity relationships and suggests that reduction of hydrophobicity within the β-sheet regions may be the primary driver of bacterial membrane selectivity (**Fig. 4**).

Previous studies using mostly alpha-helical peptides have shown insertion of polar or charged residues into the hydrophobic face of AMPs can create imperfect amphipathicity and increased membrane selectivity^23^. Highly hydrophobic or amphipathic peptides generally have a strong propensity for self-aggregation which has been shown to influence mammalian toxicity for some AMPs. Our PG-1 data also supports a relationship between highly aggregative variants and increased toxicity. Additionally, machine learning algorithms suggest that reducing overall charge and increasing hydrophobicity in the tail and loop regions, especially by mutation to tryptophan, generally increases mammalian membrane specificity. These changes would also be expected to increase the likelihood of aggregation.

One intriguing hypothesis is that PG-1 membrane selectivity is dictated by the localized concentration required to trigger self-aggregation at the cell membrane. Bacterial membranes are highly anionic, mammalian membranes less so. This would explain why PG-1.0 disrupts bacterial membranes at low concentration but mammalian membranes require a significantly higher concentration. Increasing the critical concentration required to trigger self-aggregation by reducing peptide hydrophobicity and/or amphipathicity could prevent aggregation at the lower concentrations found near mammalian membranes but still trigger self-aggregation at increased local concentrations found near highly anionic bacterial membranes. This is something we will look to further explore in future studies.

The strategy of combining dmSLAY with machine learning offers the unprecedented ability to rapidly explore AMP sequence-structure-function relationships with far greater depth than ever before. This strategy can be applied to many AMP sequences to help elucidate new ways to improve antimicrobial peptide therapeutic optimization as well as synthetic antibiotic peptide drug design.

## Materials and Methods

### Minimum inhibitory concentration assays

Stock peptides were diluted to 256 µg/ml in 350 µls of either MH media and 100 µl was aliquoted in the top row of a polypropylene 96 well plate in triplicate. Peptides were then serial diluted two-fold down columns of the plate in MH. *E. coli* W3110 or were grown overnight in LB media at 37 °C. Cells from overnight cultures were diluted to a concentration of ∼1 × 10^6^ cells/ml in MH and 50 µl were added to each well of the 96 well plate containing diluted peptide. Plates were incubated at 37 °C for 18-24 hours and examined by eye for growth. Final MIC was reported as the median of triplicate assays.

### Protegrin-1 library cloning

A library insert was first generated by PCR using forward primer oJR557 with reverse primer oJR560 and 2x(NR)tether gBlock as the template (Table S3). The insert and the pMMBEH67_lpp_ompA vector were digested with KpnI and SalI and ligated overnight at 4°C using T4 ligase. The ligated library was cleaned and transformed into *E. coli* W3110 competent cells via electroporation for further analysis via SLAY.

### Deep mutational SLAY scan

*E. coli* W3110 frozen cells containing the BH plasmid library were diluted 1:1000 and recovered in 10 ml of MH supplemented with 75 μg/ml carbenicillin for 2 hours. The culture was then back diluted to OD 0.05 and three triplicate 5 ml cultures were set up in MH supplemented with 75 μg/ml carbenicillin. Triplicate reactions included: Uninduced (0 µM IPTG), and induced (100 µM IPTG). All triplicate cultures were then grown for 4 hours at 37°C. Plasmids from each triplicate culture were then miniprepped and Illumina sequencing primers were used to produce an amplicon via PCR (Table S3). Amplicons were sent to Genewiz for next generation sequencing by Illumina MiSeq technology with 30% added Phi-X DNA.

### Deep mutational SLAY analysis

The reads obtained from next-generation sequencing underwent a meticulous series of filtering steps to isolate only the reads containing the desired amplicon regions. To achieve this, we initially utilized SeqKit^27^, a powerful tool for sequence manipulation, to specifically select sequences that matched the desired amplicon regions. This ensured that only the relevant reads were retained for further analysis. Following the initial filtering step, we applied stringent criteria to exclude reads that did not meet the required standards. Specifically, reads with a quality score below Q30 were removed from the dataset. This quality threshold ensured that only reads with a high-accuracy sequence, indicative of reliable data, were retained for downstream analysis. To remove any remaining adapter sequences, we employed Flexbar^28^ — a versatile tool for read preprocessing, which we used to trim the reads, focusing solely on the variable region of interest—the actual peptide sequence that carries the biological information we sought.

Through the application of SeqKit, and Flexbar, we successfully isolated the desired amplicon regions and obtained high-quality reads with accurate peptide sequences. These processed reads formed the foundation for subsequent analyses, where we applied built-in programs available in the Linux operating system. These programs enabled us to efficiently sort and count the unique reads, providing us information about the abundance and diversity of the peptide sequences within our dataset. The resulting counts of peptide reads were then utilized as input for DESEQ2^29^, a software package for differential gene expression analysis. By comparing the read counts between two libraries, DESEQ2 allowed us to identify peptides that exhibited significant differences in their read counts. This analysis provided us insights into the variations in peptide abundance and expression levels between the two libraries under investigation reported as log2-fold change scores.

### Peptide synthesis

All peptides used in this work were synthesized commercially by GenScript’s custom peptide synthesis service and analyzed by RP-HPLC and mass spectrometry to confirm molecular weight. For high purity peptides (> 90%) final concentration was determined using the molecular weight, A_205_, and A_205_ extinction coefficient. Final concentrations of these peptides were also adjusted for purity. A full list of all of the peptides used and their reported commercial purity can be found in supplemental data file 1.

### Hemolysis assay

Single donor human red blood cells were washed in PBS and adjusted to a concentration of 1 × 10^9^ cells/ml. Each peptide was added in triplicate to 200 μl of cells at a concentration of 128 μg/ml in a 96 well polypropylene plate. PBS alone and 1% triton-X100 were used for background normalization and 100% hemolysis respectively. Plates were incubated for 3 hours at 37 ºC. Following incubation samples were centrifuged at 800 g for 20 min and 100 μl of supernatant was transferred to a flat bottom 96 well plate. Percent hemolysis for each sample was determined by normalizing the absorbance at 540 nm for each sample to the average background and dividing by the average absorbance for 1% Triton X-100 (100% hemolysis). Error bars represent one standard deviation of triplicate samples.

### Circular dichroism spectroscopy

Stock peptides were diluted in 10 mM phosphate (pH 7.4) to 200 μg/ml in a volume of 200 μl. Samples were incubated 1-2 hours at room temperature and then analyzed using a Jasco-815 CD spectrometer with a 0.1 cm path-length quartz cuvette at the Targeted Therapeutic Drug Discovery & Development Program Core at UT Austin. The CD spectra were collected using far-UV spectra (195-250 nm) with background corrected for 10 mM phosphate buffer alone. Reported spectra are an average of three separate spectra obtained from the same sample.

### Aggregation assays

In a NUNC black-walled clear bottom 96-well place PG-1 variants were diluted to 128 µg/ml in triplicate to a volume of 98 µl in PBS. 2 µl of prepared PROTEOSTAT detection reagent was then added to each well. The plate was then mixed for 10 seconds in a Biotek Synergy LX plate reader and allowed to incubate in the dark for 10 minutes after which the fluorescence was read using the red filter cube (ex. 530/25, em. 590/35). Triplicate samples were normalized to PBS alone and reported as the mean.

### Propidium iodide uptake

Single colonies from overnight growth on LB were inoculated into MH broth and grown to mid-log phase. Cells were then washed twice with PBS + 50 mM glucose and resuspended to an OD = 0.1 in PBS + 50 mM glucose. Propidium iodide was added at a concentration of 10 µg/ml and 50 µL of the propidium iodide cell mixture was quickly added to a pre-prepared 96-well plate (NUNC black-walled clear bottom plate) that contained 50 µl of various peptide concentrations. The plate was then allowed to incubate for 30 minutes in the dark at 37°C and was then read using the Biotek Synergy LX plate reader with the fluorescence red filter cube (ex. 530/25, em. 590/35). Triplicate samples were normalized to non-treated wells and presented as a mean ± one standard deviation.

### Kill curve

*E. coli* W3110 cells were back diluted from an overnight LB culture into MH to an OD of 0.001. Peptides stock solutions were diluted in MH to 8 µg/ml. 100 µl of diluted cells were combined with 100 µl of diluted peptide stock in triplicate. 10 µl of each sample were then serially diluted ten-fold at the indicated time points and plated on LB agar. CFUs were counted after overnight growth at 30°C. Error bars represent one standard deviation of triplicate samples.

### Protein representation and transfer learning

To ensure a robust representation of the peptides, we utilized transfer learning from the esm2 15 billion parameter model^30^, a pre-trained protein Language Model (pLM), developed by Facebook. Features learned by the pLM can be transferred to any task (prediction) requiring a numerical representation (embeddings) by extracting these embeddings after the pre-training stage^31^. This method enabled us to obtain numerical embeddings for each peptide sequence, resulting in a high-quality protein representation. The embedding extraction can be found on the esm2 GitHub page (https://github.com/facebookresearch/esm#main-models). Since the extracted embeddings contained more than five thousand features, we needed to address the issue of overfitting. To mitigate this, we employed a feature selection technique to choose only those features that exhibited a high correlation with the target variable. We fine-tuned the correlation threshold to maintain a minimal set of features, approximately equivalent to the number of samples in the training set. This approach allowed us to alleviate overfitting while retaining the most relevant and informative features.

### Machine learning modeling

We have implemented a consensus machine learning model that determines whether a peptide sequence can be classified as a promising hit or a potential antimicrobial peptide. To achieve this classification, the peptide sequence must meet the cutoff thresholds of three distinct models trained to predict peptide activity (MIC), hemolysis, and selectivity. To handle undefined values in the MIC model, such as >64 μg/ml MIC, they were converted to 100 μg/ml MIC. Subsequently, peptide values with MIC ≤ 8 μg/ml were classified as active peptides, while those above this threshold were classified as non-active. These MIC-based classifications of peptides were utilized to train a classification model capable of predicting antimicrobial peptide activity based on its amino acid sequence. In the hemolysis model, we utilized empirically obtained hemolysis percentages from laboratory experiments. These experimental values served as the target to train a regression model capable of predicting the percentage of hemolysis induced by each peptide. In addition, we introduced a new target variable called ‘selectivity’ by multiplying the MIC value by the hemolysis value and subsequently applying a logarithmic transformation (log10). This new selectivity score was then used to train a regression model capable of predicting the level of selectivity for each peptide. A lower selectivity score indicates an active peptide that is also non-hemolytic.

The dataset consisted of empirical data collected from 96 PG-1 variants (Data file 1). To train and evaluate our models, we divided this dataset into a training set, which encompassed 80% of the original data selected as a random sample, and a separate test set containing the remaining 20%. We employed various algorithms, including Random Forest, Support Vector Machine (SVM), Lasso, and Decision Tree, among others, to train our models. To optimize their performance, we fine-tuned the hyperparameters of these algorithms using a Grid search from the Scikit-learn library. For each target, we selected the best-performing model based on its performance. Detailed information and implementation can be found on our GitHub page (https://github.com/ziul-bio/Protegrin-1_Slay_and_ML) in the Jupyter Notebooks 04-06.

## Acknowledgements

We would like to thank the Targeted Therapeutic Drug Discovery and Development Program at the University of Texas for access to circular dichroism training and equipment.

## Author Contributions

Conceptualization: JR, BD, LV,

CW Methodology: JR, LV

Investigation: JR, LV

Visualization: JR, LV

Funding acquisition: BD, CW

Project administration: JR, BD

Supervision: BD, CW

Writing-original draft: JR

Writing - review and editing: JR, LV, BD, CW

## Data Sharing Plan

Raw sequencing data from dmSLAY will be uploaded to the SRA database. Coding materials for sequence analysis and machine learning are available at (https://github.com/facebookresearch/esm#main-models) and (https://github.com/ziul-bio/Protegrin-1_Slay_and_ML). All other data is available in the main text or supplementary files.

