## Supplemental Figures and Tables for "Deep mutational scanning and machine learning uncover antimicrobial peptide features driving membrane selectivity"

**This PDF includes:**

Supplemental Figures S1-S5
Supplemental Tables S1-S4

A

Alanine scan of Protegrin-1.

| Name | Residue Change | MIC |
|---|---|---|
| **PG-1.0** | RGGRLCYCRRRFCVCVGR | **4** |
| PG-1 1A | A----------------- | 8 |
| PG-1 2A | -A---------------- | 8 |
| PG-1 3A | --A--------------- | 8 |
| PG-1 4A | ---A-------------- | 16 |
| PG-1 5A | ----A------------- | 8 |
| PG-1 6A | -----A------------ | 4 |
| PG-1 7A | ------A----------- | 16 |
| PG-1 8A | -------A---------- | 4 |
| PG-1 9A | --------A--------- | 8 |
| PG-1 10A | ---------A-------- | 8 |
| PG-1 11A | ----------A------- | 4 |
| PG-1 12A | -----------A------ | 2 |
| PG-1 13A | ------------A----- | 8 |
| PG-1 A14 | -------------A---- | 4 |
| PG-1 15A | --------------A--- | 8 |
| PG-1 16A | ---------------A-- | 2 |
| PG-1 17A | ----------------A- | 8 |
| PG-1 18A | -----------------A | 16 |

MIC: minimum inhibitory concentration

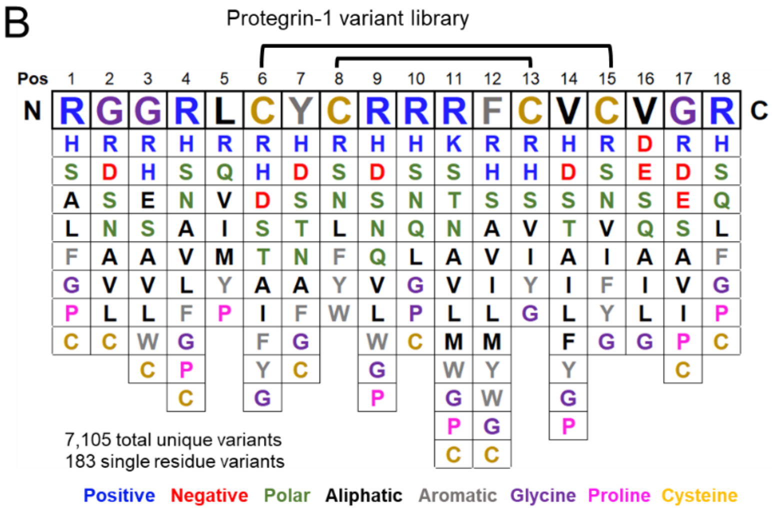


**Figure S1. Alanine scan and Protegrin-1 variant library.** A) an alanine scan performed on the native Protegrin-1 (PG-1.0) amino acid sequence showing antibacterial activity (MIC) in µg/ml. B) Chart of the variance found within the Protegrin-1 dmSLAY library. The native Protegrin-1 sequence is shown at the top with mutations observed at each location within the library below. Amino acids are color coded by side chain similarity. Brackets represent where disulfide bonds are present.

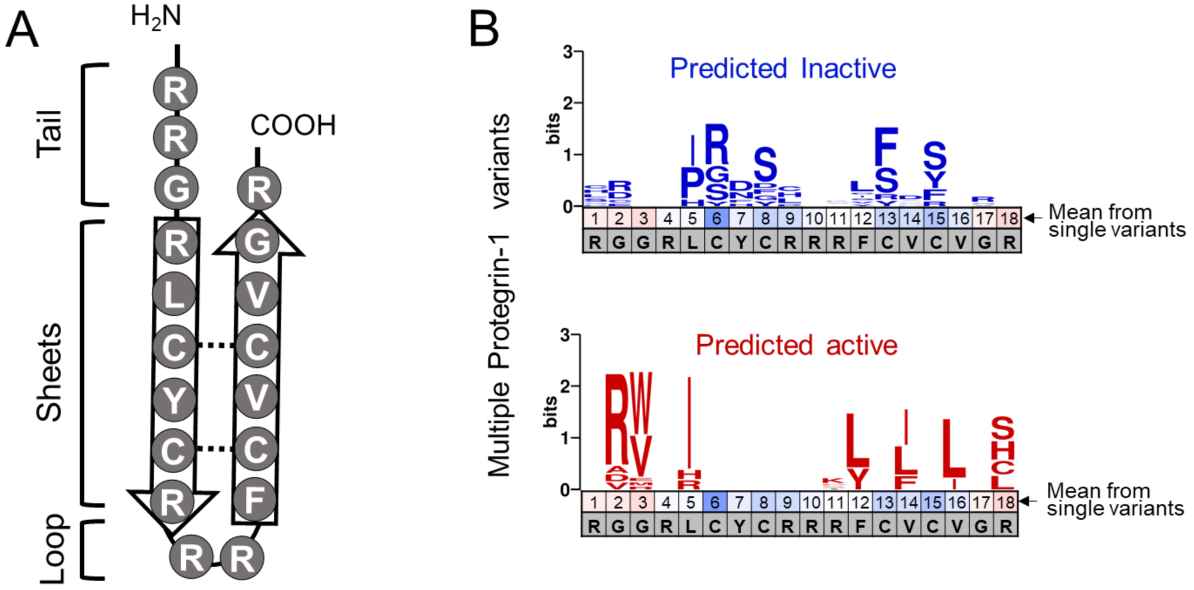


**Figure S2: Protegrin-1 structure and multiple variant dmSLAY predictions.** A) a diagram of the sequence and secondary structure found in Protegrin-1 divide into three regions: loop, sheets, and tail. beta-sheets are represented by arrows and disulfide bonds by dashed lines. B) Sequence logo plots showing mutations found in the 50 multiple Protegrin-1 dmSLAY variants with the highest (top) and lowest (bottom) $\log_2$-fold change in reads. The native Protegrin-1 sequence is shown in gray and mean average $\log_2$-fold change prediction from single residue dmSLAY variants is color coded by position.

A

dmSLAY active serine and histidine containing PG-1

| Name | Residue Change | Changes | L2FC | MIC | %Hemo | SS |
|---|---|---|---|---|---|---|
| **PG-1.0** | RGGRLCYCRRRFCVCVGR | **0** | **-1.00** | 4 | 56.4 | 226 |
| PG-1.49 | -----------Y-----H | 2 | -7.30 | 2 | 32.2 | 64 |
| PG-1.50 | -R-----F----G----H | 4 | -3.69 | 4 | 18.0 | 72 |
| PG-1.51 | -----------L-----S | 2 | -3.42 | 8 | 16.5 | 132 |
| PG-1.52 | --VH-------Y------ | 3 | -3.21 | 8 | 46.7 | 373 |
| PG-1.53 | --V--------------S | 2 | -3.18 | 64 | 24.9 | 1596 |
| PG-1.54 | -RVH-------Y------ | 4 | -3.15 | 32 | 29.2 | 933 |
| PG-1.55 | -RV--------S----Q- | 4 | -3.09 | 16 | 6.3 | 101 |
| PG-1.56 | ------FRS---S----- | 4 | -2.96 | 4 | 18.2 | 73 |
| PG-1.57 | -------------A---H | 2 | -2.92 | 2 | 21.4 | 43 |
| PG-1.58 | --V--------L-----S | 3 | -2.89 | 32 | 13.8 | 441 |

L2FC: mean $log_2$-fold change, MIC: minimum inhibitory concentration (µg/ml)
%Hemo: percent hemolysis, SS: selectivity score (MIC*%Hemo)

B

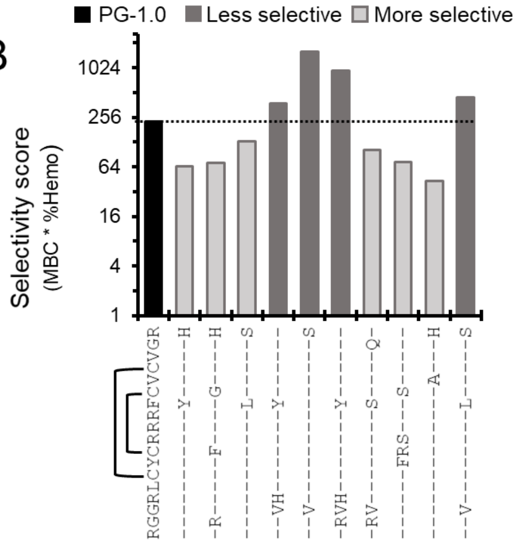


**Figure S3: Selectivity of serine and histidine containing PG-1 variants.** A) Table showing the biochemical characteristics of serine and histidine containing Protegrin-1 variants from dmSLAY. MIC is the median of triplicate assays and %Hemolysis is the mean of triplicate assays. B) Bar chart showing the selectivity score of serine and histidine containing variants on a $log_2$ scale. Residue changes are shown below. Brackets show where disulfide bonds are formed in the native structure.

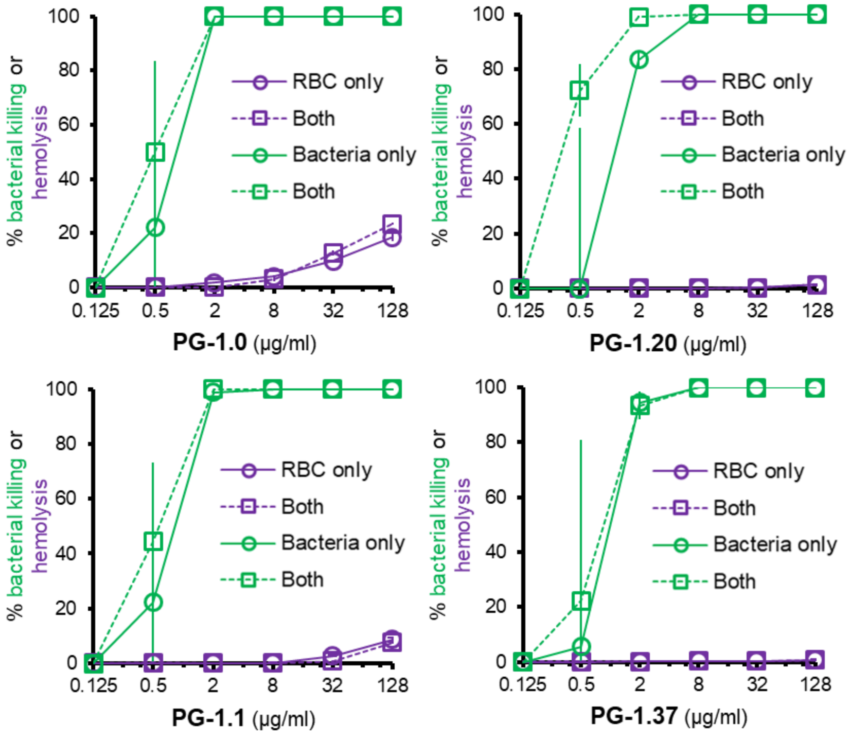


**Figure S4: Comparing Protegrin-1 variant activity in mixed cultures.** Graphs of PG-1 (top left), PG-1.1 (bottom left), PG-1.20 (top right), and PG1-37 (bottom right) percentage of bacterial killing (green) and % hemolysis (purple) with 1 x $10^9$ red blood cells (RBC), 1 x $10^6$ E. coli W3110 cells (Bacteria) or both at various concentrations shown on a $\log_2$ scale. Each data point is the mean of triplicate reactions and error bars are one standard deviation.

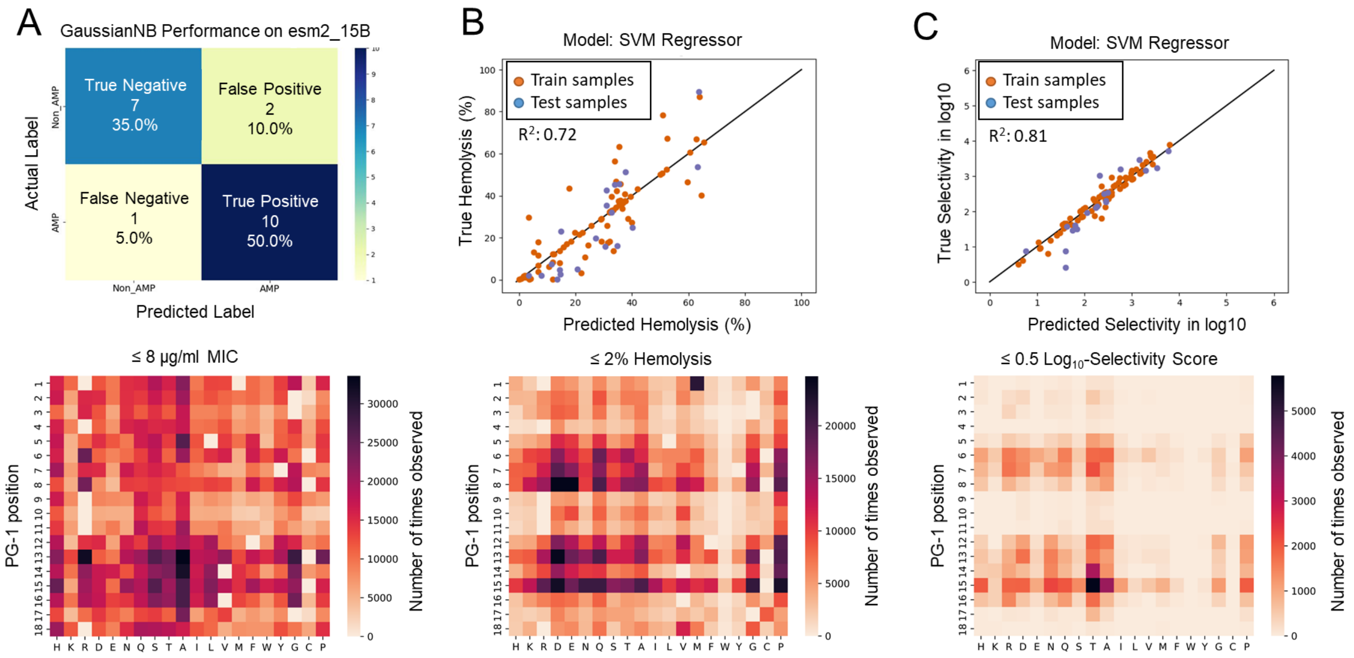


**Figure S5. Training of machine learning models and specific attribute mutational profiles.** **Top panels:** Categorization of 20% of data from a model predicting variant antibacterial activity. Non_AMP (MIC > 8 µg/ml), AMP (MIC ≤ 8 µg/ml) (A) predicted versus true hemolysis for trained (80%) and test (20%) data (B) or predicted versus true $log_{10}$-selectivity score (C). **Bottom panels**: Mutational profiles of variants from 5.7 million candidates with a predicted MIC ≤ 8 µg/ml (A) % hemolysis ≤ 2 (B) or $log_{10}$-Selectivity score ≤ 0.5.

### Table S1. dmSLAY in vitro Protegrin-1 variant data.

| Name | Sequence | Changes | L2FC | padj | MIC (µg/ml) | Mean %Hemo | SS |
|---|---|---|---|---|---|---|---|
| PG-1.0 | RGGRLCYCRRRFCVCVGR | 0 | -1.00 | 2E-76 | 4 | 56.4 | 226 |
| PG-1.1 | RGGRLCYCRRRSCVCVGR | 1 | -0.833 | 2E-47 | 8 | 5.0 | 40 |
| PG-1.9 | RGWRLCYCRRRFCVCVGR | 1 | -1.675 | 4E-178 | 16 | 89.4 | 1430 |
| PG-1.10 | RGGRLCFCRRRFCVCVGR | 1 | -1.714 | 3E-202 | 32 | 29.0 | 928 |
| PG-1.11 | RGGRLCYCRRRFCFCVGR | 1 | -1.473 | 1E-200 | 64 | 13.6 | 873 |
| PG-1.12 | RGGRLCYCRRRFCVCVGL | 1 | -1.267 | 5E-120 | 32 | 45.5 | 1456 |
| PG-1.13 | RGWRLCYCRRRFCVCVRR | 2 | -3.08 | 5E-197 | 16 | 60.7 | 971 |
| PG-1.14 | RGVRLCYCRRRLCVCVGR | 2 | -2.71 | 8E-127 | 8 | 43.5 | 348 |
| PG-1.15 | RGVRLCYCRRRFCVCLGR | 2 | -3.18 | 5E-151 | 8 | 67.2 | 537 |
| PG-1.16 | RGGRLCYCRRRFCVCLGS | 2 | -2.17 | 5E-93 | 4 | 63.3 | 253 |
| PG-1.17 | RGGRLCYCRRRFCFCVGL | 2 | -3.19 | 7E-111 | 32 | 37.4 | 1195 |
| PG-1.18 | RGGRICYCRRRYCICVGR | 3 | -2.40 | 1E-46 | 16 | 32.4 | 518 |
| PG-1.19 | RGGRLCYCRRKFCVCVRC | 3 | -1.73 | 1E-28 | 16 | 35.5 | 569 |
| PG-1.20 | RGGRLRYCHRRFCVSVGR | 3 | -1.65 | 3E-50 | 4 | 1.0 | 4 |
| PG-1.21 | RGVHLCYCRRRYCVCVGR | 3 | -3.21 | 0 | 8 | 34.4 | 275 |
| PG-1.22 | RGGRLCYCRRRYCVCVGH | 2 | -7.30 | 7E-08 | 16 | 18.1 | 290 |
| PG-1.23 | RGWRLCYCRRRFCICVGR | 2 | -3.80 | 4E-110 | 16 | 86.9 | 1391 |
| PG-1.24 | RGWRLCYCRRRFCVCVGC | 2 | -3.69 | 3E-11 | >64 | 78.3 | - |
| PG-1.25 | RGWRLCYCRRRFCVCVGH | 2 | -3.65 | 2E-45 | >64 | 46.4 | - |
| PG-1.26 | RGVRLCYCRRRFCICVGR | 2 | -3.58 | 7E-225 | 16 | 52.3 | 838 |
| PG-1.27 | RGWRLCYCRRRFCVCLGR | 2 | -3.55 | 2E-75 | 64 | 40.0 | 2562 |
| PG-1.28 | RGVRICYCRRRFCFCVGR | 3 | -7.25 | 3E-08 | 16 | 50.0 | 799 |
| PG-1.29 | RRGRICYCRRKFCVCVGR | 3 | -4.17 | 2E-10 | 16 | 19.8 | 316 |
| PG-1.30 | RGWSLCYCRRKFCVCVGR | 3 | -4.13 | 5E-07 | 32 | 65.4 | 2093 |
| PG-1.31 | RVVRLCFCRRRFCVCVGR | 3 | -3.56 | 1E-13 | 16 | 36.0 | 576 |
| PG-1.32 | HGWRLCYCRRRFCVCVGC | 3 | -3.55 | 2E-26 | >64 | 35.5 | - |
| PG-1.33 | LSGRLCYCRRRFCVCLRR | 4 | -3.21 | 3E-80 | 4 | 37.6 | 150 |
| PG-1.34 | RGWRLGYCRRRFCVSIGR | 4 | -3.17 | 1E-72 | 2 | 22.5 | 45 |
| PG-1.35 | RRVHLCYCRRRYCVCVGR | 4 | -3.15 | 4E-08 | 16 | 18.3 | 292 |
| PG-1.36 | RGGRLCYCRGRYCIRSGR | 5 | -2.90 | 1E-05 | >64 | 0.2 | - |
| PG-1.37 | HVRRLCYCRRRFCACVGS | 5 | -2.43 | 2E-11 | 1 | 2.6 | 3 |
| PG-1.38 | RRGRICYCPLRFCVCLVR | 6 | -2.07 | 0.0003 | >64 | 8.4 | - |
| PG-1.39 | RGGRLGYCRRRFCVCVGR | 1 | 0.3009 | 1E-13 | >64 | 6.7 | - |
| PG-1.40 | RGGRLCDCRRRFCVCVGR | 1 | 0.3402 | 6E-15 | >64 | 0.3 | - |
| PG-1.41 | RGGRLCYCRRRFGVCVGR | 1 | 0.1834 | 2E-05 | >64 | 0.6 | - |
| PG-1.42 | RGGRLCYCRRRFCDCVGR | 1 | 0.2873 | 9E-12 | >64 | 0.1 | - |
| PG-1.43 | RGGRLSYCLRRFCVCVGR | 2 | 0.82 | 2E-40 | >64 | 12.2 | - |
| PG-1.44 | RGGRLCYSRRRFCVCVGS | 2 | 0.79 | 5E-23 | >64 | 15.6 | - |
| PG-1.45 | RGGRLCYCHRRFFVRVGR | 3 | 1.17 | 2E-50 | 32 | 2.0 | 63 |
| PG-1.46 | RGRRLCYCHRRFCDSVGR | 4 | 1.35 | 0.0004 | >64 | 0.1 | - |
| PG-1.47 | GRERLCYCHRRLCVCVGR | 5 | 1.46 | 3E-21 | >64 | 2.0 | - |

L2FC: $\log_2$-fold change in reads, padj: adjusted p value, MIC: minimum inhibitory concentration
%Hemo: percent hemolysis, SS: selectivity sccore (MIC * %Hemo)

**Table S2. Bacterially selective machine learning PG-1 variants.**

| Name | Sequence | MIC (μg/ml) | %Hemo | SS | $Log_{10}$ SS |
|---|---|---|---|---|---|
| PG-1.0 | RGGRLCYCRRRFCVCVGR | 4 | 56.4 | 226.0 | 2.35 |
| bsPG-1.1 | RGGRL**Q**YCRRRFCV**AR**GR | 16 | 0.43 | 6.9 | 0.84 |
| bsPG-1.2 | RGGRL**Q**YCRRR**G**CV**T**VGR | 4 | 0.34 | 1.4 | 0.13 |
| bsPG-1.3 | RGGRL**A**YCRRR**D**C**T**CVGR | >64 | -0.17 | - | - |
| bsPG-1.4 | RGGRL**D**YCRRRFCV**TT**GR | 64 | 0.17 | 10.6 | 1.03 |
| bsPG-1.5 | RGGRL**QA**CRRRFCV**T**VGR | 8 | 0.72 | 5.8 | 0.76 |
| bsPG-1.6 | RGGRL**R**YCRRRF**D**V**S**VGR | 16 | 0.74 | 11.8 | 1.07 |
| bsPG-1.7 | RGGRLC**TA**RRRFCV**R**VGR | 2 | 0.91 | 1.8 | 0.26 |
| bsPG-1.8 | RGGRL**T**YCRRRFC**TA**VGR | 16 | 0.04 | 0.7 | -0.15 |
| bsPG-1.9 | RGGRL**T**YCRRR**D**CV**A**VGR | 32 | 0.12 | 3.7 | 0.57 |
| bsPG-1.10 | RGGRL**A**YCRRRFCV**DT**GR | 64 | 0.09 | 5.9 | 0.77 |
| bsPG-1.11 | RGGRL**R**YCRRR**GT**VCVGR | >64 | 0.27 | - | - |
| bsPG-1.12 | RGGRL**Q**YCRRRFC**AR**VGR | 16 | 0.14 | 2.3 | 0.35 |
| bsPG-1.13 | RGGRLCYCRRR**N**CV**TT**GR | >64 | -0.06 | - | - |
| bsPG-1.14 | RGGRLC**TA**RRRFCV**H**VGR | 16 | 0.64 | 10.2 | 1.01 |
| bsPG-1.15 | RGGRLC**D**CRRRFC**NM**VGR | >64 | -0.10 | - | - |
| bsPG-1.16 | RGGRL**Q**YCRRRF**DT**CVGR | >64 | 0.03 | - | - |
| bsPG-1.17 | RGGRLC**T**CRRRFC**AS**VGR | 32 | 0.49 | 15.6 | 1.19 |
| bsPG-1.18 | RGGRL**R**YCRRRF**R**V**S**VGR | 16 | 1.53 | 24.4 | 1.39 |
| bsPG-1.19 | RGGRL**E**YCRRRFCV**NT**GR | >64 | 1.28 | - | - |
| bsPG-1.20 | RGGRL**T**YCRRRFC**AD**VGR | 64 | 0.25 | 15.9 | 1.20 |
| bsPG-1.21 | RGGRLCY**R**RRR**TA**VCVGR | 8 | 0.40 | 3.2 | 0.51 |
| bsPG-1.22 | RGGRLCYCRRRFC**TP**V**T**R | 64 | 0.39 | 25.0 | 1.40 |
| bsPG-1.23 | R**T**GRLCYCRRR**D**C**T**CVGR | 32 | 0.05 | 1.7 | 0.23 |
| bsPG-1.24 | RGGRL**H**Y**R**RRRFCV**R**VGR | 8 | 9.72 | 77.8 | 1.89 |
| bsPG-1.25 | RGGR**P**CYCRRRFC**GT**VGR | >64 | -0.01 | - | - |
| bsPG-1.26 | RGGRLC**D**CRRR**T**C**R**CVGR | >64 | 0.19 | - | - |
| bsPG-1.27 | RGGR**DI**YCRRRFCV**T**VGR | 8 | 0.66 | 5.3 | 0.72 |
| bsPG-1.28 | RGGRLCYCRRR**T**CV**DT**GR | 32 | 0.16 | 5.1 | 0.71 |
| bsPG-1.29 | RGGRLCY**S**RRRFC**KT**VGR | 64 | 0.56 | 35.9 | 1.55 |
| bsPG-1.30 | RGGRL**A**YCRRRFCV**A**V**A**R | 2 | 1.08 | 2.2 | 0.33 |
| bsPG-1.31 | RGGRLCY**G**RRRFCV**NQ**GR | 32 | 2.06 | 66.0 | 1.82 |
| bsPG-1.32 | RGGR**S**C**H**CRRRFCV**I**VGR | 4 | 15.70 | 62.8 | 1.80 |
| bsPG-1.33 | RG**R**R**A**CYCRRRFCV**H**VGR | 4 | 3.66 | 14.7 | 1.17 |
| bsPG-1.34 | RGGRL**S**YCRRRFCV**AG**GR | 16 | 0.32 | 5.1 | 0.71 |
| bsPG-1.35 | RG**Q**RLCYCRRRF**N**VC**T**GR | 32 | 0.40 | 12.8 | 1.11 |
| bsPG-1.36 | RGGRLCYCRRRFC**AVN**GR | 64 | 0.12 | 7.5 | 0.87 |

bsPG: bacterially selective Protegrin-1 machiine learning variant, MIC: minimum inhibitory concentration, %Hemo: percent hemolysis,SS: selectivity score (MIC * %Hemo)
**bold**: residue change

**Table S3. Evaluation of machine learning performance.**

| Group | n | Improved SS* | Average $log_2$-fold change in SS* | MIC ≤ 8 | Hemolysis < 2% | $Log_{10}SS$ < 0.5 |
|---|---|---|---|---|---|---|
| Bacterially specific ML | 36 | 81% | -3.76 | 25% | 89% | 17% |
| dmSLAY active | 31 | 16% | 1.63 | 29% | 6% | 3% |

n: sample number, SS: selectivity score (MIC * %hemolysis), MIC: minimum inhibitory concentration, ML: machine learning
* relative to PG-1.0

**Table S4. Plasmids, Strains, and Oligonucleotides.**

| Plasmids | Source |
|---|---|
| pMMBEH67_lpp_ompA | (16) |
| **Strains** | **Source** |
| *E. coli* W3110 | Lab Stock |
| **Oligonucleotides** | **Sequence** |
| oJR557 - F dmSLAY library | gtattggtaccagtcaagagcctg |
| oJR560 - R Protegrin-1 dmSLAY primer | CTG CAG GTC GAC TTA (N1:94020202)(N2:02940202)(N3:02029402) (N4:02020294)(N2)(N2) (N4)(N1)(N2) R(N2)(N1) (N1)(N1)(N2) R(N2)(N1) (N3)(N1)(N1) (N2)(N2)(N4) (N3)(N2)(N3) (N1)(N2)(N3) R(N2)(N1) R(N4)(N1) R(N2)(N1) (N1)(N1)(N3) (N1)(N2)(N3) (N2)(N2)(N2) (N1)(N2)(N2) (N3)(N2)(N3) GGT TCC TCC GAT ACC CGC AG |
| 2x(NR)tether gBlock | ATTGCCGATGGTACACGTCAAGTCAAGAGCCTGCAGCGCCCGCCGCAG AGGCGACTCCTGCTGCTGAAGCTCCAGCTAGCGAAGCGCCTGCAGCAG AAGCTGCCCCAGCGGATGCTGCCGAAGCCCCAGCCGCTGGCATCAGTC AGGAACCTGCTGCACCAGCTGCGGAAGCTACACCAGCAGCGGAGGCAC CAGCGAGTGAAGCACCGGCTGCGGAAGCCGCTCCTGCAGATGCCGCT GAGGCTCCAGCTGCGGGTATCGGAGGAACCCGCGGTGGGCGTCTTTGT TA |
| F amplicon | aatgATACGGCGACCACCGAGATCTACACTCTTTCCCTACACGACGCTCT TCCGATCTCTCCAGCTGCGGGTATCGGAGGA |
| R index 1 | CAAGCAGAAGACGGCATACGAGATCGTGATGTGACTGGAGTTCAGACG TGTGCTCTTCCGATCTgccaagcttgcatgcctgcaggtcgacTTA |
| R index 2 | CAAGCAGAAGACGGCATACGAGATACATCGGTGACTGGAGTTCAGACG TGTGCTCTTCCGATCTgccaagcttgcatgcctgcaggtcgacTTA |
| R index 3 | CAAGCAGAAGACGGCATACGAGATGCCTAAGTGACTGGAGTTCAGACG TGTGCTCTTCCGATCTgccaagcttgcatgcctgcaggtcgacTTA |
| R index 4 | CAAGCAGAAGACGGCATACGAGATTGGTCAGTGACTGGAGTTCAGACG TGTGCTCTTCCGATCTgccaagcttgcatgcctgcaggtcgacTTA |
| R index 5 | CAAGCAGAAGACGGCATACGAGATCACTGTGTGACTGGAGTTCAGACGT GTGCTCTTCCGATCTgccaagcttgcatgcctgcaggtcgacTTA |
| R index 6 | CAAGCAGAAGACGGCATACGAGATATTGGCGTGACTGGAGTTCAGACG TGTGCTCTTCCGATCTgccaagcttgcatgcctgcaggtcgacTTA |
| R index 7 | CAAGCAGAAGACGGCATACGAGATGATCTGGTGACTGGAGTTCAGACG TGTGCTCTTCCGATCTgccaagcttgcatgcctgcaggtcgacTTA |
| R index 8 | CAAGCAGAAGACGGCATACGAGATTCAAGTGTGACTGGAGTTCAGACGT GTGCTCTTCCGATCTgccaagcttgcatgcctgcaggtcgacTTA |
| R index 9 | CAAGCAGAAGACGGCATACGAGATCTGATCGTGACTGGAGTTCAGACGT GTGCTCTTCCGATCTgccaagcttgcatgcctgcaggtcgacTTA |
| R index 10 | CAAGCAGAAGACGGCATACGAGATAAGCTAGTGACTGGAGTTCAGACGT GTGCTCTTCCGATCTgccaagcttgcatgcctgcaggtcgacTTA |
| R index 11 | CAAGCAGAAGACGGCATACGAGATGTAGCCGTGACTGGAGTTCAGACG TGTGCTCTTCCGATCTgccaagcttgcatgcctgcaggtcgacTTA |
| R index 12 | CAAGCAGAAGACGGCATACGAGATTACAAGGTGACTGGAGTTCAGACGT GTGCTCTTCCGATCTgccaagcttgcatgcctgcaggtcgacTTA |

All oligonucleotides were ordered from Integrated DNA technologies (IDT). N1, N2, N3 and N4 are hand mixed %bases (i.e. N1:94020202 = N1 is a mixture of 94% A, 2% C, 2% G, 2% T)
